# Evolutionary trajectory of receptor binding specificity and promiscuity of the spike protein of SARS-CoV-2

**DOI:** 10.1101/2022.06.11.495733

**Authors:** Cyril Planchais, Alejandra Reyes-Ruiz, Robin Lacombe, Alessandra Zarantonello, Maxime Lecerf, Margot Revel, Lubka T. Roumenina, Boris P. Atanasov, Hugo Mouquet, Jordan D. Dimitrov

## Abstract

SARS-CoV-2 infects cells by attachment to its receptor – the angiotensin converting enzyme 2 (ACE2). Regardless of the wealth of structural data, little is known about the physicochemical mechanism of interactions of the viral spike (S) protein with ACE2 and how this mechanism has evolved during the pandemic. Here, we applied experimental and computational approaches to characterize the molecular interaction of S proteins from SARS-CoV-2 variants of concern (VOC). Data on kinetics, activation- and equilibrium thermodynamics of binding of the receptor binding domain (RBD) from VOC with ACE2 as well as data from computational protein electrostatics revealed a profound remodeling of the physicochemical characteristics of the interaction during the evolution. Thus, as compared to RBDs from Wuhan strain and other VOC, Omicron RBD presented as a unique protein in terms of conformational dynamics and types of non-covalent forces driving the complex formation with ACE2. Viral evolution resulted in a restriction of the RBD structural dynamics, and a shift to a major role of polar forces for ACE2 binding. Further, we investigated how the reshaping of the physicochemical characteristics of interaction affect the binding specificity of S proteins. Data from various binding assays revealed that SARS-CoV-2 Wuhan and Omicron RBDs manifest capacity for promiscuous recognition of unrelated human proteins, but they harbor distinct reactivity patterns. These findings might contribute for mechanistic understanding of the viral tropism, and capacity to evade immune responses during evolution.

## Introduction

Coronaviruses are enveloped viruses with single-stranded RNA genome. Seven members of this family are known to infect humans, all causing respiratory infections, albeit with different severity ^1^. Severe acute respiratory syndrome coronavirus 2 (SARS-CoV-2) was first detected in the late 2019 and caused a pandemic with tremendous societal consequences. Since its identification the virus has mutated and a number of divergent variants emerged^2^. Some of these variants have markedly increased transmissibility and ability to evade humoral immunity induced by vaccination or previous infection. Such SARS-CoV-2 variants are classified by the WHO as Variants of Concern (VOC). The increased transmissibility and evasion of antibody recognition by SARS-CoV-2 is driven by mutations in spike (S) protein^2^. The S protein is a trimer made of identical protomers, each consisting of S1 and S2 subunits. S1 subunit is responsible for virus attachment to the host cells, whereas S2 subunit mediates penetration of the virus into host cells ^3, 4^. The S1 subunit contains a receptor binding domain (RBD) responsible for interaction with the receptor for SARS-CoV-2, the angiotensin converting enzyme 2 (ACE2) ^5^. Mutations acquired during the evolution of SARS-CoV-2 can also occur in the RBD. For instance, SARS-CoV-2 Alpha variant (B.1.1.7) has a single amino acid replacement in the RBD (N501Y), whereas Omicron variant BA.1 (B.1.1.529) has 15 RBD mutations ^6^. Notably, the mutations in the RBD are often located at the molecular surface directly contacting ACE2. The mutations in the RBD can reshape the binding surface, and increase the binding affinity of S for ACE2, which ultimately contribute to enhanced infectivity of SARS-CoV-2 variants ^7–9^.

Characterizing the molecular mechanism of SARS-CoV-2 S – ACE2 interaction is paramount for understanding how the virus infects host cells, and how it can evade immune recognition. Up to now, there is a large body of comprehensive structural information from cryo-electron microscopy and X-ray crystallography studies describing some molecular details of the interaction of ACE2 with S (or isolated RBD domains) from the original Wuhan strain and from different VOCs ^4, 10–17^. Atomic level structures of the S protein in complex with neutralizing antibodies have also been solved and contributed to understanding the mechanisms of viral neutralization ^15, 16, 18, 19^. These structural analyses clearly demonstrated that the receptor binding site of RBD was remodeled by introduction of amino acid replacements during the evolution of SARS-CoV-2. However, these analyses did not depict the functional consequences of mutations in the RBD. Indeed, the structural analyses provide only a partial portrayal of the mechanism of intermolecular interaction. They describe static final states of the interacting proteins but cannot directly quantify the energetic changes occurring during different phases of intramolecular interaction. The dissection of binding energies characterizing the interaction process may provide information about the structural dynamics and the types of non-covalent binding forces driving complex formation ^20–23^. During the ongoing pandemic there has been a sequential emergence of different SARS-CoV-2 variants with increased infectivity and capacity for evading immune recognition. It is not understood how this evolution trajectory correlates with changes in the receptor binding mechanism of S protein. Moreover, it is not known how the viral evolution influence the binding of S protein to unrelated to ACE2 human proteins (off-targets binding).

In the present study, we investigated the mechanism of recognition of ACE2 by the RBD. We used a biosensor technology to experimentally evaluate activation- and equilibrium thermodynamics of the interaction of the RBD of original Wuhan strain of SARS-CoV-2 and of VOCs – Alpha (B.1.1.7), Beta (B.1.351), Gamma (P.1), Delta (B.1.617.2) and Omicron (BA.1; B.1.1.529). We performed both experimental and computational analyses to assess the type of non-covalent interactions driving the molecular recognition of ACE2 by SARS-CoV-2. We also evaluated the functional repercussions of accumulating mutations in S proteins in terms of ACE2 specificity and promiscuous binding to unrelated proteins. Our findings revealed profound reshaping of the molecular interactions driving the binding of ACE2 by RBDs from different SARS-CoV-2 variants. This reconfiguration resulted in changes in off-target reactivities (binding promiscuity) of S proteins of SARS-CoV-2 – an important finding contributing for better understanding the mechanism of virus infectivity, tropism, and the evasion of immune recognition.

## Results

### Evolution of the receptor binding domain of S protein of SARS-CoV-2

Since its identification in late 2019, SARS-CoV-2 has constantly evolved ^2, 6^. Certain mutations of the virus, especially those affecting the S protein, can result in increased transmission, virulence and ability to evade antibody responses ^2^. The virus strains harboring such mutations are classified as VOCs. Owing to its superior fitness, SARS-CoV-2 variant Omicron (B.1.1.529, and its subvariants) has now largely replaced the other VOCs, and it is responsible for most of the present COVID-19 infections worldwide ^8^. Mutations that are localized in the RBD of S protein can alter the physicochemical features of binding surface that is involved in interaction with ACE2. Indeed, depicting electrostatic charges and amino acid residues involved in RBD-ACE2 contacts demonstrate considerable alterations in the receptor binding surface in different VOCs (**Fig. 1A**). These changes were not associated, however, with a considerable difference in the size of the binding footprint area (**Fig. 1B**). Previous studies have reported that mutations in the receptor binding motif (RBM) can increase the binding affinity for ACE2, thus contributing to the enhanced infectivity of SARS-CoV-2 ^13, 16, 19, 24, 25^. Accordingly, RBD and S proteins from Wuhan and Omicron variants demonstrated marked qualitative differences in their real time binding profiles to ACE2 (**Fig. 1C**). These differences in the binding profiles translated into quantitative differences in binding kinetics and affinity of RBDs for ACE2 (see next section and **Fig. 2B**). Collectively, these data suggest that the changes in the physicochemical characteristics of the receptor binding site on RBD correlate with changes in the mechanism of molecular recognition of ACE2. Since alternation in the recognition mechanism can explain important traits of SARS-CoV-2 VOCs such as better infectivity and evasion of neutralization by antibodies, we focused our efforts to decipher the evolutionary trajectory of the physicochemical characteristics of RBD-ACE2 binding mechanisms.

**Figure 1.**
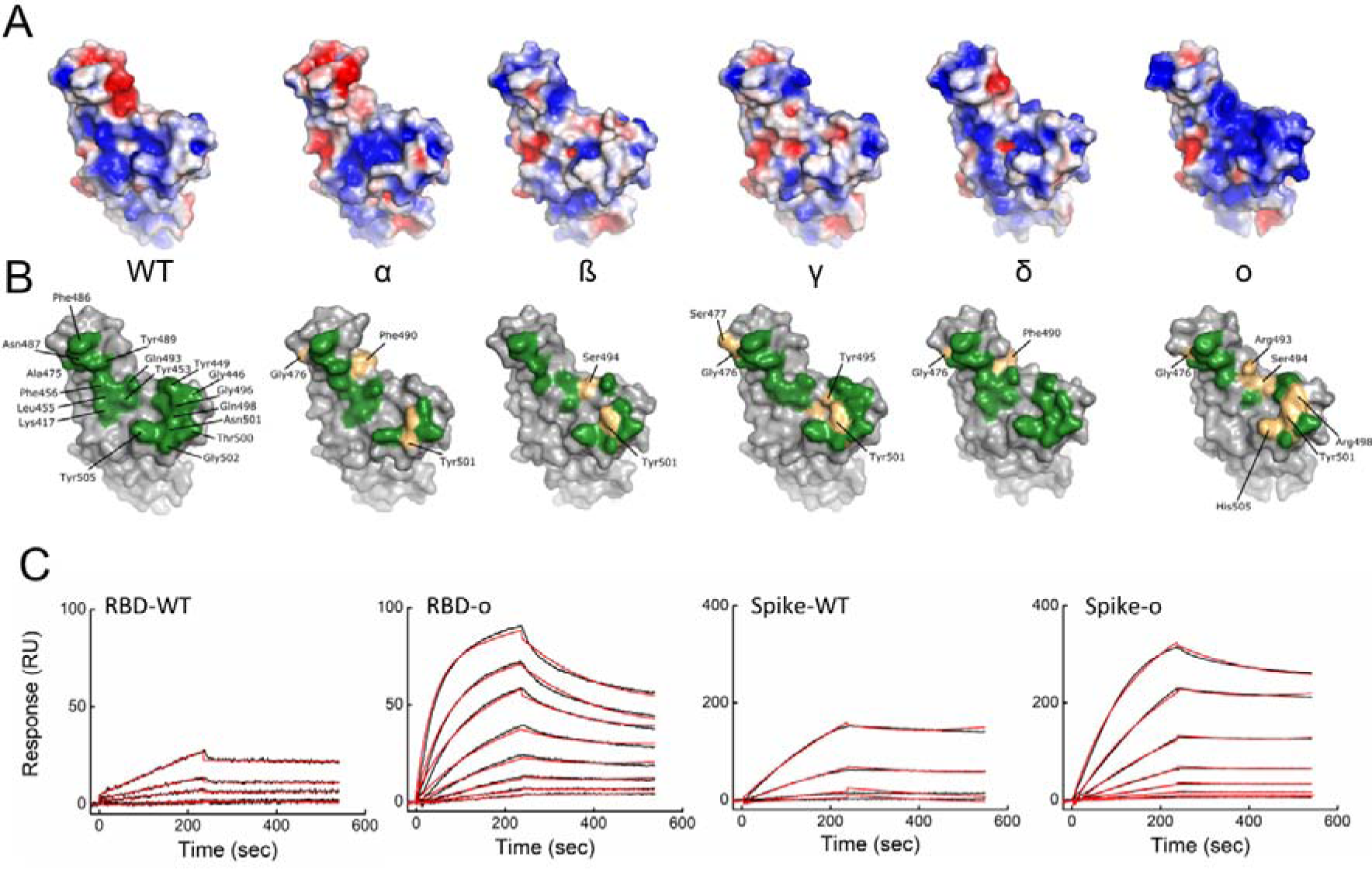
Evolution of the receptor binding site of RBD of S protein of SARS-CoV-2. (A) Depiction of the electrostatic charges of the binding site for ACE2 on RBD from WT and different VOCs of SARS-CoV-2. Blue – positive charges; Red – negative charges. For depiction of the electrostatic charges PyMOL molecular graphics software was used. (B) Binding footprint of ACE2 on the surface of RBD. The contact residues are indicated in green and displayed for RBD-Wuhan. The implication of novel residues (due to mutation or conformational changes) is indicated in gold. The models of RBDs were generated using following PDB files 6LZG (Wuhan, indicated on graph as WT); 7EDJ (B.1.1.7 or α variant); 7V80 (B.1.351 or β variant); 7NXC (P.1 or γ variant); 7W9I (B.1.617.2 or δ variant), and 7T9L (B.1.1.529 or o variant). (C) Real-time interaction profiles of the binding of RBD-Wuhan and RBD-o (left panels) and S-Wuhan and S-o (right panels) to sensor-chip immobilized human ACE2. The RBDs, and S trimers were injected at two-fold dilutions with concentrations starting at 15.625 and 1.25 nM, respectively. The experimental binding curves are shown in black, the curves obtained by global kinetic analysis are shown in red. All measurement were performed at 24 °C.

**Figure 2.**
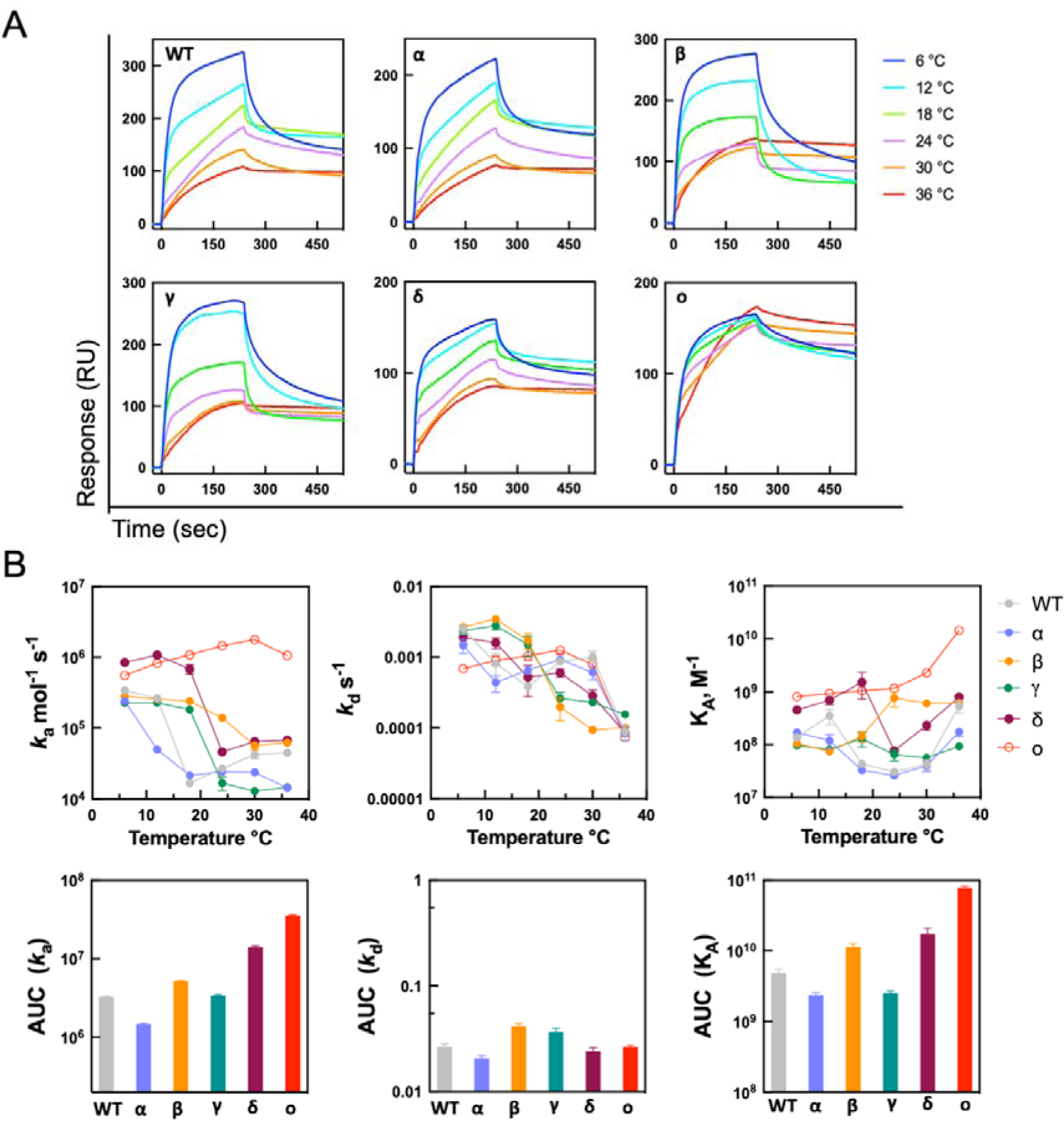
Temperature dependence of the interaction of the RBD of S protein of SARS-CoV-2 with ACE2. (A) Real-time interaction profiles of binding of RBDs from Wuhan (WT) and different VOCs with immobilized human ACE2. The binding profiles obtained at increasing temperatures: 6, 12, 18, 24, 30 and 36 °C are depicted in different colors. The binding curves were generated after injection of RBD at following concentrations: 125 nM (Wuhan and δ), 200 nM (α, β, and γ), and 62.5 nM (o). (B) Upper panels show temperature dependence of the kinetic rate constants and the equilibrium association constant of binding of different RBD strains to human ACE2. Lower panels depict the integrated areas of the corresponding curves (AUC). Each point represents the average value of the kinetic constant ±SD obtained after three independent evaluations of binding curves acquired at increasing temperatures 6, 12, 18, 24, 30 and 36 °C. Association rate constants were evaluated by global fit analysis of real time binding curves from at least 9 dilutions of each RBD variant. Dissociation rate constants were evaluated by separate kinetic fit analysis. The equilibrium association constants were calculated by the ratio *k*_a_/*k*_d_.

### Effect of temperature on the kinetics of interaction of RBD from VOCs with ACE2

Analysis of the impact of temperature on the kinetics of protein-protein interactions can provide important information about the binding process. Thus, the response of kinetics to changes in the temperature reflects the presence of conformational changes in interacting partners, solvent effects, and type of non-covalent forces driving formation of the complex. A previous study demonstrated that the kinetics and affinity of the RBD-Wuhan and the RBD-α are very sensitive to variation in the temperature ^26^. This sensitivity also reflected in differences in the capacity of SARS-CoV-2 to infect target cells – thus, virus attachment and infectivity was significantly augmented at low temperature (4 °C).

In the present study, comparing the binding profiles of RBD from the Wuhan strain and different VOCs revealed clear differences in the temperature sensitivity of their interactions with ACE2 (**Fig. 2A**). Thus, real-time profiles depicting association and dissociation phases of the binding process showed that the interaction of RBD-Wuhan with ACE2 was characterized with high temperature sensitivity. The binding response progressively decreased as a function of the temperature. This decrease was accompanied by changes in the shape of the binding curves indicating that the temperature strongly influences the binding kinetics of the RBD-Wuhan. Interaction profiles of RBDs from VOCs (α, β, γ, δ) also demonstrated high sensitivity to temperature, albeit it had a different impact on the binding profiles of various variants (**Fig. 2A**). Contrary to all the other variants, changes in temperature resulted in a qualitatively distinct effect on the interaction of the RBD of Omicron (o) variant with ACE2 (**Fig. 2A**). Thus, the variation of the temperature in the range 6 - 36 °C had only a negligible effect on the binding intensity. Notably, RBD-ο reached the greatest intensity of binding to ACE2 at the highest temperature – an effect opposite to what was observed with RBDs from other variants (**Fig. 2A**). Taken together, these data suggest that the RBD-ο uses a qualitatively different molecular mechanism for binding to ACE2 as compared to Wuhan strain and other VOCs.

To provide a deeper understanding of these differences and obtain quantitative data, we next evaluated the binding kinetics and equilibrium affinity for the interactions of SARS-CoV-2 Wuhan and VOC RBDs with ACE2 (**Fig. 2B**, upper panels). The obtained results demonstrated that the temperature has a strong influence on the association rate constant (*k*_a_) of all RBD variants. However, the *k*_a_ for RBD-o presented a qualitatively distinct behaviour compared to the other variants. Thus, with the sole exception of RBD-ο, an increase of the interaction temperature resulted in a marked reduction of the *k*_a_ values, even though this decrease had unique profiles for RBDs from different variants of SARS-CoV-2 (**Fig. 2B**, upper panels). In contrast, the *k*_a_ that characterizes the binding of RBD-o to ACE2 was generally augmented with increased temperature. In addition, the interaction of RBD-o with ACE2 was characterized by a very rapid association (*k*_a_ ca. 10^6^ mol^-1^ s^-1^) across the entire temperature range. The fast association rate and robustness of *k*_a_ of RBD-o to temperature variation in the whole range was apparent when comparing the areas under the curves of the association rates of different RBDs (**Fig. 2B**, lower panels). In contrast to the heterogenous effects of the temperature on the association rate constant, the *k*_d_ values for the interactions of different RBDs with ACE2 were influenced more uniformly, as evidenced by the kinetics plots and the area under the curves (**Fig. 2B**). With increase of the temperature, there was generally a decrease in the *k*_d_ i.e., the stability of the complexes of RBDs with ACE2 increased. Finally, when comparing the effects of temperature on the binding affinity (equilibrium association constant, K_A_) for ACE2, it was also obvious that K_A_ of RBD-o manifested the most distinct behavior. Collectively, these data strongly suggest that evolution of SARS-CoV-2 resulted in qualitative and quantitative changes in the molecular mechanism driving the interaction with ACE2. Especially, these data strongly imply that the RBD from Omicron variant uses a different molecular mechanism for binding to its receptor.

### Activation and equilibrium thermodynamics analyses reveal divergent molecular mechanism for RBD Omicron binding to ACE2

The temperature dependencies of the kinetic rate constants and equilibrium affinity constant contain quantitative data on the binding process ^27, 28^. To acquire this information and uncover the energetic changes that are taking place during the interaction of the RBD with ACE2, we applied the approach of Arrhenius for evaluation of the activation thermodynamics. Figure 3A depicts the Arrhenius plots, which represent the natural logarithm of the rate constants versus reciprocal temperature (in Kelvin degrees). The figure also shows Van’t Hoff plot that characterizes the temperature dependency of equilibrium association constant. The Arrhenius plots for the association phase clearly highlighted the distinct response of RBD-o as compared to other variants observed in Figure 2. The Arrhenius plots allow calculation of activation energies characterizing the association and the dissociation, and further of the changes of the activation thermodynamic parameters. These analyses revealed that interaction with ACE2 of RBD from SARS-CoV-2 Wuhan and from VOCs (α, β, γ, δ) were characterized with identical pattern of changes in the thermodynamic parameters during association. Thus, as observable on Figure 3B, all bindings were accompanied by large negative values of the activation entropy (TΔS^‡^ in the range from ∼ -90 to -135 kJ mol^-1^, for different VOC), and negative values in enthalpy (ΔH^‡^ in the range from ∼ -45 to -90 kJ mol^-1^, for different VOC). Notably, qualitatively different patterns of the energetic changes were observed in the case of the association of the RBD Omicron variant (**Fig. 3B**). The extend of changes in activation entropy were significantly reduced as compared to all other variants (TΔS^‡^ = -21 ±8.2 kJ mol^-1^). Strikingly, in contrast to all other variants, the association of RBD-o to ACE2 was accompanied by changes in enthalpy with positive value (ΔH^‡^ = 17.7±7.2 kJ mol^-1^). The activation thermodynamic changes describing the dissociation of RBD from ACE2 followed similar pattern for the Wuhan strain and all VOCs (**Fig. 3B**). Both enthalpy and entropy changes presented a negative sign, although with a certain difference in absolute values. For instance, the dissociation of RBD-β from ACE2 was characterized by a ΔH^‡^ of -102.8 ±20.6 kJ mol^-1^, as compared with ΔH^‡^ of -54.6 ±26.2 and -40 ±29.6 kJ mol^-1^ for RBD-Wuhan and RBD-o, respectively. This difference could explain the observation that changes in the thermodynamic parameters at equilibrium had a very similar pattern for the interactions of RBD-β and RBD-o (**Fig. 3B**). Nevertheless, the pathways for reaching this similar pattern at equilibrium were qualitatively different for the two variants.

**Figure 3.**
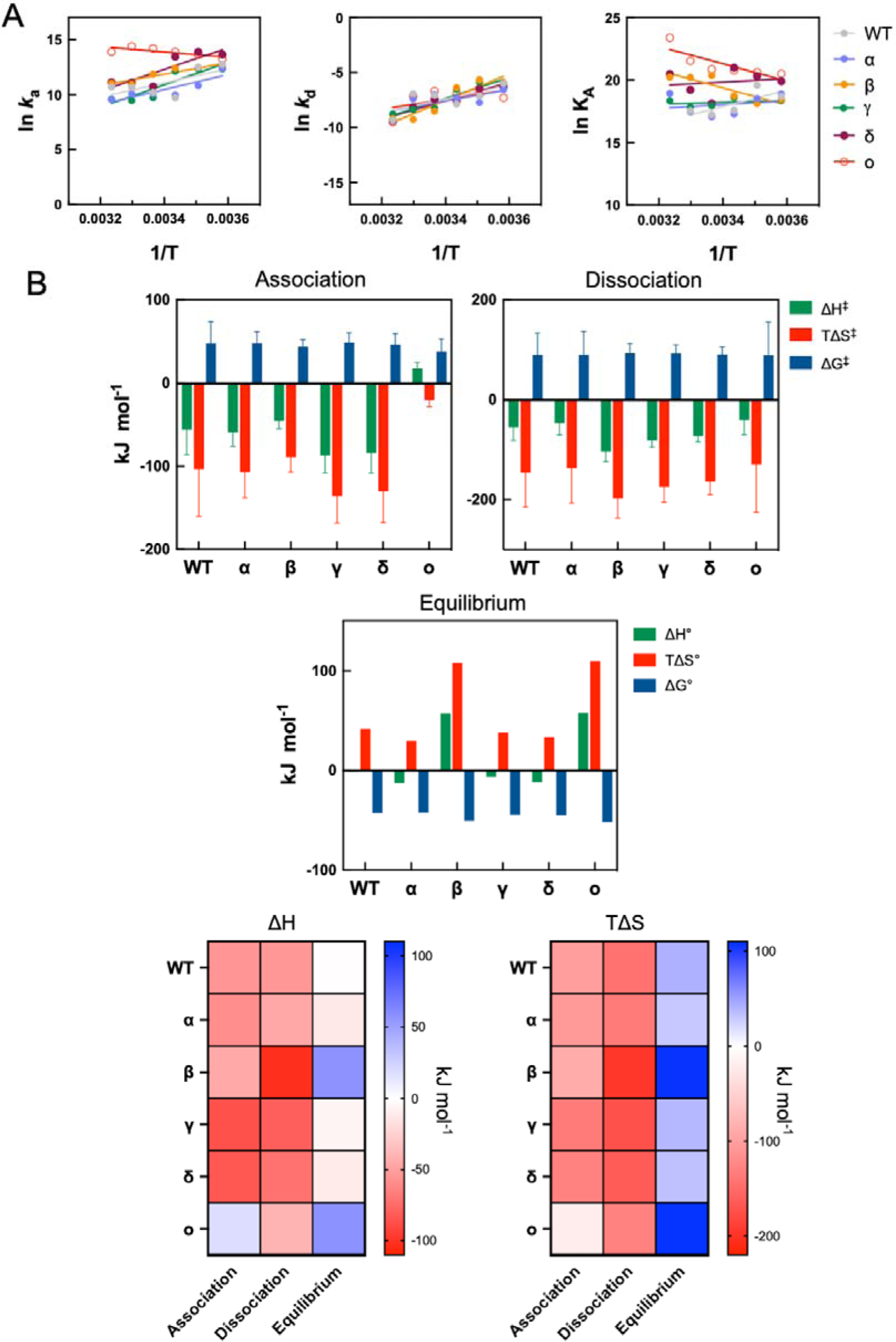
Activation and equilibrium thermodynamics of binding of different variants of RBD to ACE2. (A) Arrhenius plots (left and central panel) and van’t Hoff plot (right panel) depicting the temperature dependence of kinetic rate constants and equilibrium association constant, respectively of binding of RBDs from different strains of SARS-CoV-2 to ACE2. The linear regression analysis was used for determination of the slopes of the plots. (B) Changes in the activation and equilibrium thermodynamic parameters (enthalpy, entropy, and Gibbs free energy) of interaction of different variants of RBD with ACE2. Heat maps present overview of the pattern of changes of the entropy and enthalpy for different phases of the binding process. All thermodynamic parameters were evaluated at reference temperature of 24 °C or 297.15 K.

Taken together, activation and equilibrium thermodynamic data revealed that the evolution of SARS-CoV-2 was accompanied by a profound resetting of the molecular mechanism of binding to ACE2. In contrast to the RBD Wuhan and other VOCs, RBD-o uses a qualitatively different mechanism of recognition of its ligand. The thermodynamic changes, strongly suggest that this variant associates with ACE2 with substantially diminished penalty due to structural dynamics (evident by the reduced negative value of TΔS^‡^) and by using different types of non-covalent forces driving the association phase (evident by the quantitively and qualitative change in the value of ΔH^‡^).

### Analyses of non-covalent forces driving the interaction of RBD with ACE2

The changes of enthalpy during the association phase depends on the type of the non-covalent forces driving the formation of intermolecular complex ^20, 23, 28^. Since the association ΔH^‡^ of RBD-o had a qualitatively distinct signature than those of the Wuhan strain and other VOCs, we focused our efforts to comprehend the origin of this difference. First, we performed, pH-dependence analyses of the binding of RBDs from different SARS-CoV-2 variants to immobilized ACE2 (**Fig. 4A**). The binding of RBDs to ACE2 was characterized by a clear pH-dependency – described by bell-shaped binding curves. Although interaction of RBD of Wuhan strain and VOCs (α, β, γ, δ) with ACE2 manifest similar pH dependency, RBD-o was characterized by a considerable increased sensitivity to pH in the acidic range (**Fig. 4A**). All RBDs from SARS-CoV-2 variants had similar behavior in the basic region of pH curve (i.e. pH ≥ 7.5). The high sensitivity to acidic pH of interaction of RBD-o strongly implies that ionic interactions have an important contribution for stabilization of the complex with ACE2. This result is concordant with the observed increased area of positive charges on the receptor binding site of RBD-o (**Fig. 1A**) ^29^. To gain further insight into the role of non-covalent forces driving the interactions of SARS-CoV-2 with its ligand, we evaluated the binding kinetics and the affinity as a function of the salt concentration (**Fig. 4B**). The association rate constant of binding of RBDs from SARS-CoV-2 variants (Wuhan, α, γ, δ) to ACE2 significantly decreased when the NaCl concentration of the interaction buffer was increased from 75 to 150 mM. Further increase to 300 mM had a little impact on the rate constant. Interestingly, the association rate of RBD-β was barely reduced with salt concentrations ranging from 75 to 150 mM, but a significant reduction in the association rate was observed when the binding was studied in buffer containing 300 mM NaCl (**Fig. 4B**). The most peculiar behavior of association rate as a function of salt concentration was observed in the binding of RBD-o to ACE2. Thus, increase of NaCl concentration from 75 to 150 mM resulted in a substantial augmentation (∼10-fold) of *k*_a_. Further increase of the NaCl concentration to 300 mM resulted in reduction of the rate constant. The observed discrepancies in the sensitivity of association of RBD to ACE2 to the effect of concentration of ions confirms that evolution of SARS-CoV-2 is accompanied by substantial alteration of the type of non-covalent forces driving association with the ligand.

**Figure 4.**
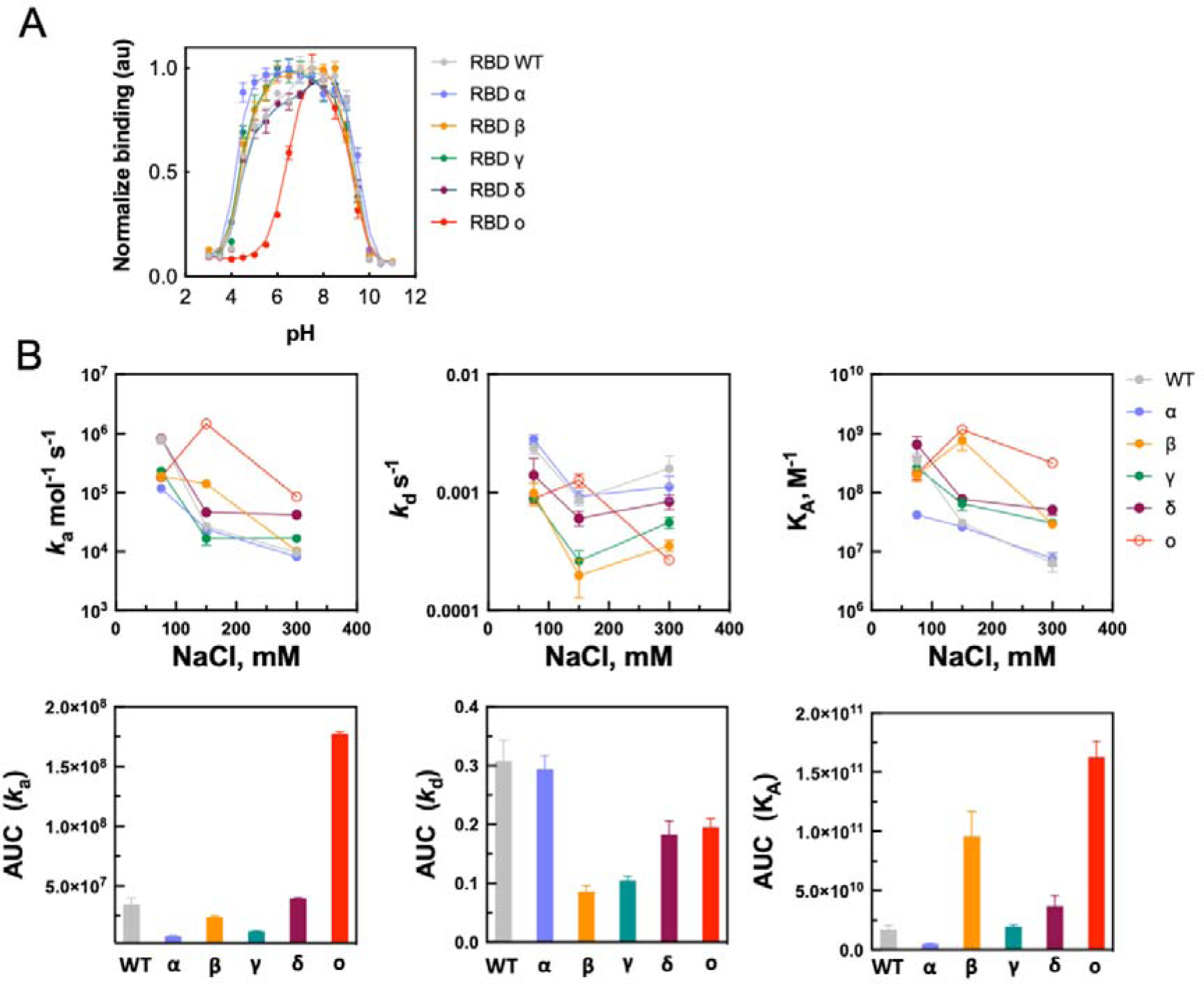
Effect of pH and ionic concentration on the binding of RBDs from different SARS-CoV-2 variants to ACE2. (A) ELISA – binding of RBD from different strains of SARS-CoV-2 with ACE2. Site-specifically biotinylated RBDs were diluted to 0.625 µg/mL, in buffers with increasing pH (0.5 units of increment of each point) and incubated with surface immobilized human ACE2. Each point represents the normalized binding intensity obtained from three repetition. A representative result out of three independent repetitions is shown. (B) Effect of ionic concentration on the binding kinetics of RBD from different variants of SARS-CoV-2 with ACE2. Upper panels show the effect of variation in the ionic concentration on the kinetic rate constants and the binding affinity. Lower panels show the integrated area of the corresponding curves (AUC). The kinetic measurements were performed by surface plasmon resonance in 10 mM HEPES buffer, containing 75, 150 or 300 mM NaCl. Association rate constants were evaluated by global fit analyses of real time binding curves from at least 9 dilutions of each RBD variant. Dissociation rate constant were evaluated by separate kinetic fit analyses. The equilibrium association constants were calculated by the ratio *k*_a_/*k*_d_. All measurements were performed at 24 °C.

Distinct effect of changes in the salt concentration were also observed in the case of dissociation rate constant of RBD-o. Thus, in contrast to all other variants, an increase of NaCl concentration from 150 mM to 300 mM resulted in considerable drop in the dissociation rate of RBD-o from ACE2 (**Fig. 4B**). Other variants demonstrate a prominent reduction in the *k*_d_ when the concentration of NaCl was increased from 75 to 150 mM. In this range *k*_d_ of RBD-o showed an opposite behavior. The changes in the concentration of ions in the buffer differentially impacted the binding affinity as well. Interestingly, the K_A_ characterizing the binding of RBD-o and RBD-β manifested similar pattern of sensitivity of the buffer salt concentration (**Fig. 4B**). Thus, there was an increase in the affinity when the NaCl concentration was increased from 75 to 150 mM. All other variants demonstrated diminished affinity for ACE2 in this condition. Importantly, the similar trend in the sensitivity of K_A_ of RBD-o and RBD-β to salt concentration is in accordance with identical changes in the enthalpy at equilibrium observed for these variants of SARS-CoV-2.

Finally computational analyses of protein electrostatics of RBDs from different VOC corroborated the experimental data. Thus, the RBD of Omicron variant demonstrated the most distinct behavior as compared to the other variants in terms of the pH dependence of the total free energy (**Fig. 5A**); the electrostatic component of the free energy (**Fig. 5B**); net charge (**Fig. 5C**), as well the extend of variation in the values of the scalars of the electric moments (**Fig. 5D**). The effect of the viral evolution on the electric moments of the RBD can be visualized alternatively by displaying electric moment vectors. The figure comparing the pH sensitivity of the electric movement vectors of RBD Wuhan and RBD Omicron clearly showed dramatic reconfiguration of this electrostatic parameter (**Fig. 5E**).

**Figure 5.**
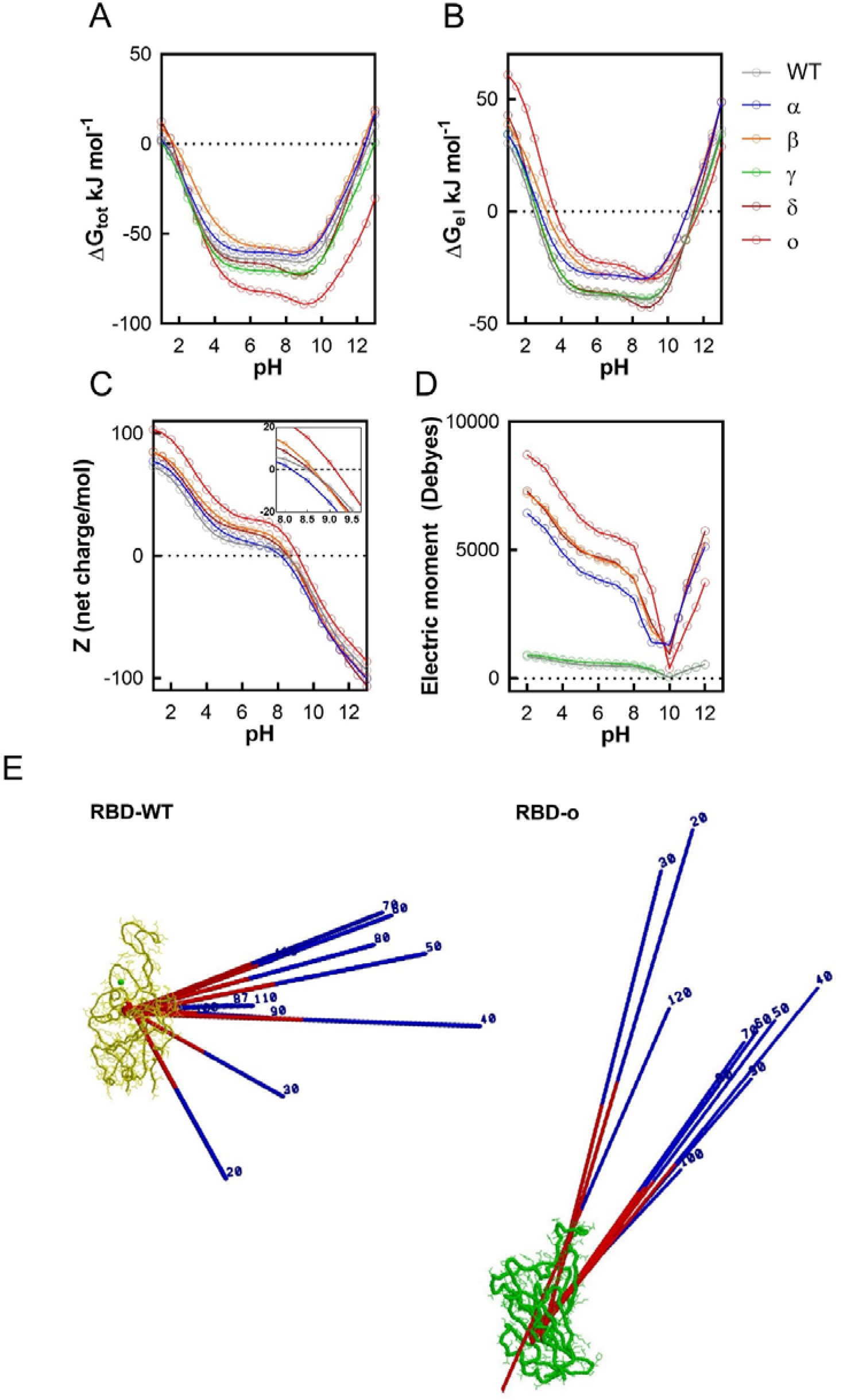
Computational electrostatics analyses of RBD from Wuhan strain and different VOC. (A) Theoretical analysis of the pH-dependence of total free energies [ΔG_tot_ (pH)] of RBDs. (B) Theoretical analysis of the pH-dependence of the electrostatic term of the free energy [ΔG_El_ (pH)] of RBDs. (C) Theoretically predicted potentiometric titration Z (pH) curves of RBDs. The inset shows magnified curves in the region of the isoelectric points of the studied proteins. Of note the titration curve of Z (pH) of RBD-γ is not presented as its calculation was not possible. (D) Theoretically predicted scalar values of the electric moments (µ) of RBDs as a function of pH. (E) Theoretically predicted vectors of electric moments (µ) of RBD Wuhan (in yellow) and RBD Omicron (in green) as function of pH. The predicted direction and size of electric moment at different pH is shown on the diagram. Please note that the structural models of both proteins are not on same scale. Computational analyses of the protein electrostatics were performed by using following PDB files 6LZG (Wuhan, indicated on graph as WT); 7EDJ (α variant); 7V80 (β variant); 7NXC (γ variant); 7W9I (δ variant), and 7T9L (o variant).

Taken together the data from pH dependence and salt concentration dependence analyses revealed that evolution of SARS-CoV-2 is accompanied by changes in the nature of non-covalent forces that contribute for association with ACE2 with the most dramatic difference observed in case of RBD-o.

### Analyses of binding promiscuity of S protein and RBD of SARS-CoV-2

Previous studies demonstrated that the S protein of SARS-CoV-2 can establish interactions with other targets unrelated to ACE2 via its RBD motif (reviewed in ^30^). The considerable reconfiguration of the physicochemical mechanism driving the interaction of the RBD with ACE2 as result of evolution of SARS-CoV-2, prompted us to evaluate the tendency for off-target interactions i.e., binding promiscuity, of the most distinct variants of the virus. First, by using ELISA we elucidated the binding of S trimer from SARS-CoV-2 Wuhan and Omicron strains with an arbitrarily selected panel of 17 target biomolecules, which consisted predominantly by proteins but also included LPS and chondroitin sulfate (**Fig. 6A**). The S proteins were able to recognize off-targets albeit with considerably lower intensity as compared with binding to the principal receptor for the virus, ACE2 (**Fig. 6A**). Thus, the S proteins demonstrated a moderate binding promiscuity as evident by reactivity to cytochrome C, nerve growth factor-beta, histone 3 and LysM (*Enterococcus faecalis*). Interestingly, both S proteins demonstrated differences in the binding intensity to distinct off-target molecules. Further, we tested the capacity of proteins present in calf fetal serum (lacking antibodies against S protein) to inhibit binding of S proteins to ACE2 and different off-targeted proteins (**Fig. 6B**). The presence of serum proteins did not affect the interaction of S proteins to ACE2. However, the excess of protein present in the serum strongly inhibited all promiscuous reactivities of S protein. Notably, a different sensitivity to inhibition by serum proteins was observed for S proteins of Wuhan strain and Omicron variant and this sensitivity varied for different target proteins (**Fig. 6B**). For both S trimers the interaction with histone 3 was most strongly affected by the presence of excess of serum proteins. Thus, even at 512-folds dilution of fetal calf serum the binding of S proteins to histone 3 was only approximately 30 % of the interaction observed in buffer only (**Fig. 6B**).

**Figure 6.**
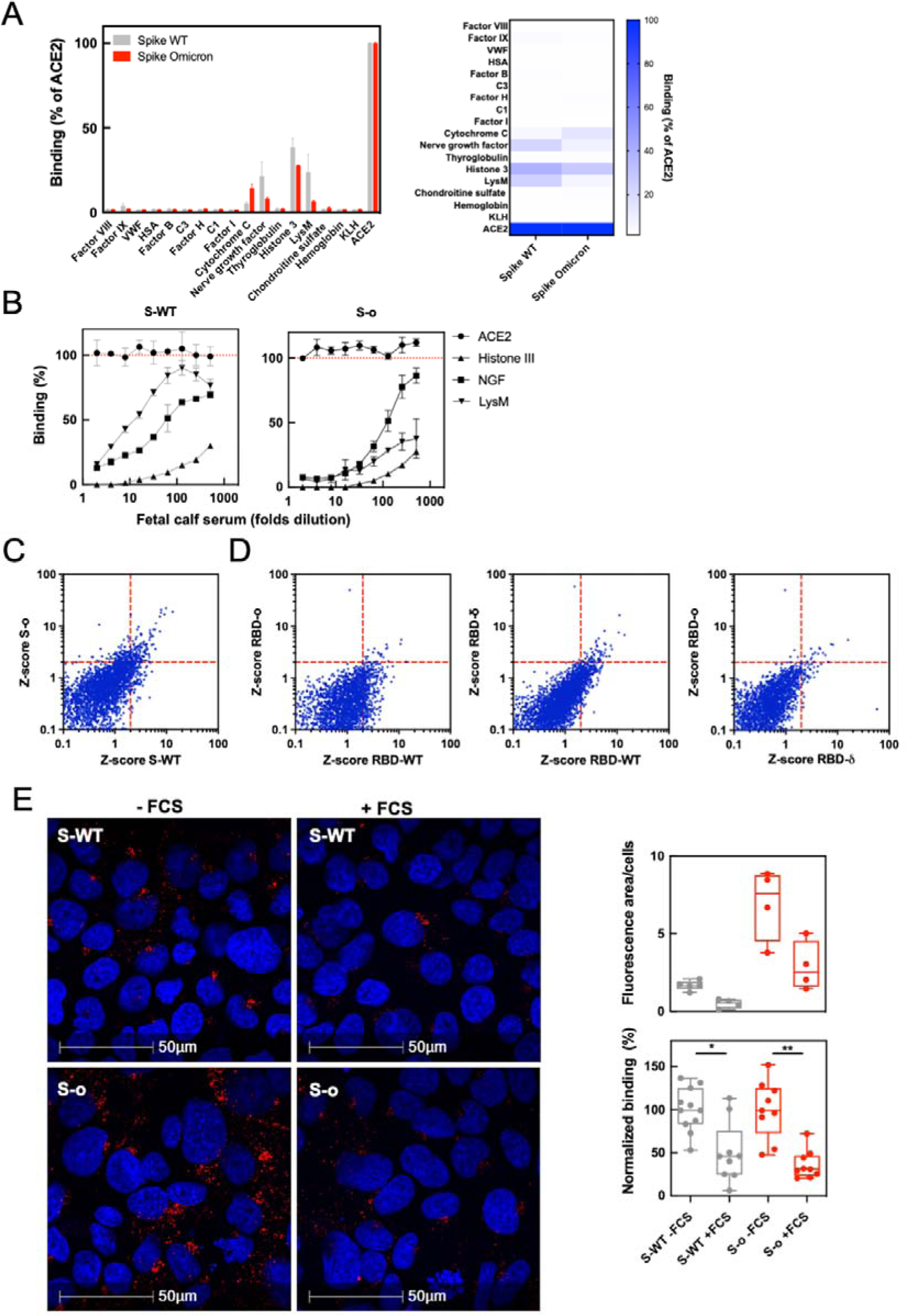
Analyses of binding promiscuity of S protein of SARS-CoV-2. (A) ELISA – reactivity of S proteins from Wuhan and Omicron to a panel of distinct proteins, and chondroitin sulfate. The graph shows the percentage binding of biotinylated S proteins (Wuhan and Omicron variants) to various target molecules as compared to their reactivity to ACE2 (considered as 100 %). The data are presented also as a heat map (right panel). Each bar represents average binding percentage obtained from five replicates in two independent experiments. (B) ELISA – reactivity of S proteins to immobilized human ACE2 and off-targeted proteins in the presence of fetal calf serum. Biotinylated S proteins were diluted in buffer containing different concentrations of FCS (starting from 50 %, followed by two-fold dilutions). The reactivity of S proteins to the respective protein in the absence of serum is shown as 100 % and indicated on the graph as the red dashed line. Each point represents average binding obtained from two replicates. (C and D) Human protein array analyses. The plots show the binding of S proteins (C) or indicated RBDs (D) to large panel of human proteins. The S trimers and RBDs were covalently labeled with Alexa Fluor 555. The binding intensity is represented as Z score – reactivity to each human protein is shown as a blue circle. Thresholds of significance i.e., 2 × Z score (corresponding to p = 0.05) are shown as red dashed lines. (E) Confocal microscopy analyses of binding of S trimers to BeWo cells. Fetal calf serum inhibits the interaction of AlexaFluor555-labelled S protein of Wuhan or Omicron variants of SARS-CoV-2 with BeWo cells. Representative images from two independent experiments showing interaction of S proteins (red) with the BeWo cells, captured by confocal microscopy. Nuclei are stained in blue. The right panels show quantification of fluorescence results from one experiment (each dot represents independent reading) or from two independent experiments after normalization – 100 % represents the average reactivity of S proteins in the absence of FCS. One-way Anova, Kruskal-Wallis-parametric analysis, with multiple comparisons, * *p* = 0.05; ** *p* = 0.01. Further details about quantification of the fluorescence signal from confocal microscopy are provided in *Material and Methods*.

To obtain a better understanding about the binding promiscuity of S proteins and about how evolution impacted this feature, we applied protein microarray technology. This approach allowed to follow the interaction of fluorochrome-labeled S proteins or RBDs with >9000 human proteins (**Fig. 6C**). The obtained results indicated that S proteins of Wuhan strain and Omicron variant manifested an overall similar extend of promiscuous binding to human proteins. However, certain proteins were preferentially recognized by one of the variants. S trimer is a large protein and it may contact off-targets with different domains apart of the RBD. Moreover, RBD binding surface for ACE2 in S proteins can be not constantly exposed due to tendency of trimer to exist more frequently in a closed state, in which the RBD molecular surfaces are not solvent exposed. Therefore, we elucidated the reactivity to human proteins of fluorochrome-labeled RBDs from Wuhan, Delta and Omicron variants. These analyses suggested that all RBDs are promiscuous, but RBD-Wuhan was prone to higher capacity for off-target interactions with human proteins as compared to RBD-δ and RBD-o (**Fig. 6D**).

Finally, to understand how the binding promiscuity of S proteins of Wuhan strain and Omicron variant impacts the recognition of cellular targets, we used a cellular model. BeWo is a human cell line derived from human placenta. These cells express moderate levels of ACE2 and are weakly pеrmissive to viral replication ^31, 32^. We assayed the behavior of different S proteins, selecting the variants with the most distinct in the physicochemical features. Thus, incubation of BeWo cells with fluorescently labeled-S proteins from SARS-CoV-2 Wuhan and Omicron in serum-free medium resulted in significant binding of both S proteins to the cells, as evidenced by results from confocal microscopy (**Fig. 6E**). The fluorescent area stained by S-o was larger, which might reflect the higher binding affinity of this variant for ACE2. To evaluate the extend of promiscuous reactivity, we also studied the reactivity of S proteins in the presence of 1 % of fetal calf serum, which was used as a source of excess of unrelated proteins for blocking the promiscuous interactions. The incubation of S proteins in the presence of serum resulted in a significant reduction of the binding to cells (**Fig. 6E**) – an observation indicating that a significant part of the reactivity of both S proteins is due to promiscuous bindings to different cellular targets. Importantly the proportion of the reduced reactivity in the presence of serum was equal for both S-WT and S-o, suggesting that they have similar degree of off-target binding to BeWo cells.

Collectively, these data suggested that S proteins have tendency for moderate binding promiscuity. These interactions might be reconfigured in result of alternations of the physicochemical characteristics of the binding surface of the RBD due to accumulation of mutations.

## Discussion

In the current study we applied experimental and computational approaches to analyze the physicochemical mechanism of the interaction of S protein of SARS-CoV-2 with its receptor ACE2. We elucidated the evolutionary trajectory of this mechanism by analyzing interactions of mutants of the virus that presently or historically represented variants of concern. The principal finding of this work is that the evolution of SARS-CoV-2 resulted in profound resetting of the mechanism of recognition of ACE2 by RBD of S protein. Thus, RBD of Omicron variant demonstrated considerable qualitative and quantitative differences in kinetics as well as in activation and equilibrium thermodynamics of binding to ACE2, as compared to RBDs of other strains of the virus. These disparities in the energetic changes of binding suggest that RBD-o behaves as a completely different protein in relation to RBDs of the Wuhan strain or other VOCs, a finding that cannot be perceived by using merely structural analyses. Remarkably, a similar extend of differences in the thermodynamic signature of binding as the one observed in the present study have been previously reported when interaction mechanisms of diverse categories of proteins were compared ^33–35^.

Association thermodynamics provides important information about the molecular events occurring during formation of intramolecular complexes. Our data demonstrate that binding of RBD from Wuhan and VOCs α, β, γ, and δ is accompanied by favorable changes for binding affinity in enthalpy (ΔH^‡^ with negative values) and unfavorable changes in entropy (TΔS^‡^ with negative values). The observed highly negative values of the association entropy (ca. -100 kJ mol^-1^) implies that the formation of the complex of ACE2 with RBD is associated with a substantial reduction of disorder in the system. Such reduction can be a consequence of restriction of the intrinsic structural dynamics of polypeptide chains of RBD upon binding to ACE2. Contrary to other VOCs, the association of RBD-o was characterized with only negligible changes in entropy – a fact indicating low degree of structural changes (in the interacting proteins or in the surrounding solvent) during formation of the intramolecular complex ^22, 28^. Rigidification of the binding surface of the RBD from Omicron variant could also be inferred from the kinetics data, where the association rate of only this variant was benefited by an increase of the ambient temperature. Previous studies demonstrated that the protein-protein interactions accompanied by conformational adaptations are generally negatively impacted by an increase of the temperature ^36^. In contrast, the association of well-fitting proteins that possess pre-formed rigid binding sites is favored by elevated temperatures due to rise in the kinetic energy of the system, which favors the formation of encounter complexes. Thus, experimental analyses of the association kinetics and thermodynamics suggests that evolution of SARS-CoV-2 is accompanied by a marked reduction in the molecular dynamics of the RBD. This finding is in accordance with several *in silico* studies showing that RBD-o decreases its conformational dynamics as compared to RBD-Wuhan and other VOCs ^9, 37–39^.

In addition to the considerable alteration in association entropy, the analyses of non-equilibrium thermodynamics of RBD-ACE2 interaction revealed a pronounced impact of the virus evolution on the association enthalpy. Thus, contrary to values of enthalpy determined for RBD-Wuhan and other VOCs, the association of RBD-o to ACE2 was accompanied by ΔH^‡^ with a positive sign. The changes in association enthalpy depend on the types, quantity, and energies of the non-covalent forces formed between interacting molecules ^20, 28, 40^. Therefore, opposite signs in the ΔH^‡^ implies that the evolution of SARS-CoV-2 have significantly remodeled the energies and type of non-covalent forces driving formation of complex with ACE2. Our experimental data dedicated on analyses of binding forces implicated in ACE2 recognition by the RBD supported the result from thermodynamic analyses and showed a qualitative shift in the type of non-covalent interactions in case of RBD-o. The computation analyses further corroborated these experimental data and unraveled that the electrostatic forces have the highest favorable contribution to the free energy of binding in case of the interaction of RBD-o ^24, 29, 37^.

This remodeling in the binding mechanism could well be explained by the available structural data which have demonstrated that as result of accumulation of mutations the binding surface of RBD-o increases significantly its overall positive charge (Fig. 1A) ^15, 17, 19^. Its noteworthy that results from *in vitro* evolution analyses demonstrated that a RBD variant with markedly augmented affinity for ACE2 (ca. 1000 folds) had also evident enlargement surface of positive electrostatic charges in the binding site for ACE2 ^41^. These results suggest that the tendency of both natural and *in vitro* evolution for optimization of binding to ACE2 of the spike protein is associated with extension of the positive charges in the binding surface of RBD. Such extension might improve charge complementarity with ACE2, which is acidic protein, but also might assist long distance electrostatic steering interaction to ACE2 or to cell membrane, which bears overall negative charge.

Our data indicated that one of the consequences of the remodeling of the non-covalent forces of the RBD binding to ACE2 is that this interaction with Omicron variant is characterized by a considerably enhanced sensitivity to acidic pH. This effect might be explained by ionization of histidine residues at the binding surface. Indeed, the inflex point of pH curve overlap with pKa value of His residue. The shape of pH dependence curve (i.e. the sharp increase in the binding with change in pH) implied that the effect is mediated due to ionization of a single His residue. It should be noted that one of the mutations in the ligand binding surface of RBD-o is replacement of Tyr505 with His (Y505H). The structural analyses demonstrate that this His establishes contacts with amino acids from ACE2 ^15, 17, 19^. This His residue might be an important contributor for overall energy of binding, and it may control the interaction with ACE2. Histidine residues are known to mediate pH sensitivity of other protein-protein interactions in acidic pH interval. As an example the pH dependency of the interaction of IgG with neonatal Fc receptor (FcRn) is controlled by His residues present in Fc portion of immunoglobulin molecule ^42^.

The pH has been already demonstrated to have important impact on S protein of SARS-CoV-2 ^43^. It can affect equilibrium of “open-” and “closed-” conformation state of the RBD domains in the trimer. The pH-induced transition to “closed” state was proposed to contribute to evasion of antibody responses ^43^. Our study suggests that the ionization of the binding surface of the RBD in SARS-CoV-2 VOC Omicron due to introduction of His may provide yet another pH dependent mechanism for control the interaction of S with ACE2. This effect might have contribution to the pathogenicity of the virus. Indeed, endocytic vesicles have acidic pH ∼6, a value of pH which is able to disrupt RBD-o:ACE2 interaction or eventually the interactions of RBD-o with other viral receptors / co-receptors, and hence determine the tropism and pathogenicity of the virus ^44^.

Analyses of the mechanism of molecular recognition of ACE2 by RBD exposed another trait that appeared during evolution of SARS-CoV-2. Thus, the results from analyses of equilibrium changes in the thermodynamic parameters demonstrate that Omicron variant had a qualitatively different profile in comparison to most of the studied strains (Fig 3B). Notably, ACE2 binding by RBD-β manifested an identical profile of energetic changes at equilibrium. Thus, in contrast to all other VOC, the interactions of RBD-β and RBD-o with ACE2 were characterized by an increase in the equilibrium enthalpy (ΔH° > 0). Although changes in equilibrium entropy for the bindings of all RBDs had qualitatively identical sign (in all cases TΔS° > 0), the bindings of RBD-β and RBD-o to ACE2 were characterized by substantially higher values (2-3 folds) of this parameter (Fig. 3B). Previous studies used molecular dynamics simulations for determination of the total binding energies (sum of the energies of different types of non-covalent interactions) of interactions of RBDs from VOC with ACE2 ^24, 38^. Notably, the binding of all RBDs with ACE2 were characterized with total binding energies with negative values. As computationally derived total binding energy could be considered as a proxy to ΔH°, we noted a contradiction between experimental data obtained in this study and the previous computational data. These differences could be explained by the effects of buffer system used and by the experimental setting i.e., ACE2 is surface immobilized and RBDs are in solution.

The increase of equilibrium entropy for bindings of RBDs to ACE2 could be interpreted in the light of statistical thermodynamics. In statistical physics the entropy can be represented by Bolzmann equation (S° = k_B_lnW) which describes the relationship between the thermodynamic parameter and the number of microstates available in the system (represented by quantity of W). In this work, the identified experimental values of ΔS° > 0 show considerable increases in the number of the molecular microstates in the reaction system at equilibrium, especially in the cases of the binding of RBD-β and RBD-o to ACE2. This increase can be explained by structural heterogeneity and/or heterogeneity in the structure of the solvent.

The data from activation thermodynamics indicate that pathways for achieving this identical thermodynamic profile at equilibrium, however, are distinct in both strains. This observation also clearly established that only integrative analyses of both activation and equilibrium thermodynamics can fully depict the mechanism of protein-protein interaction. RBDs of Beta and Omicron variants share a common mutation – K417N. However, at present it is difficult to explain how this common mutation could result in a similar thermodynamic profile at equilibrium. A possible explanation might be related to epistatic effects of the mutations as recently described in ^45^. It is noteworthy, that affinity (i.e. K_A_ constant for binding to ACE2) of both RBD-o and RBD-β demonstrated also similar behavior as a function of changes in the ionic concentration of the interaction buffer (Fig. 4B). Common equilibrium thermodynamic profiles and sensitivity to ion concentration of RBD-β and RBD-o during interaction with ACE2 is an intriguing observation in light of the fact that these two strains are characterized with the highest degree of resistance to neutralization by antibodies, induced by infection or by vaccination ^46^. This observation suggests that evaluation of distinctness of thermodynamic profiles can be used as a proxy for assessment of capacity of mutants of SARS-CoV-2 to evade antibody responses.

In addition to its main ligand, the S protein of SARS-CoV-2 has been demonstrated to interact with a relatively large set of unrelated molecular targets ^30, 47–56^. Notably, this binding promiscuity of S protein was shown to have important functional repercussion affecting the cell tropism and infectivity of the virus ^47, 50, 53, 54^. A large part of identified off-targets of S protein are recognized by RBD. In the present study we elucidated the impact of the virus evolution and changes in the physicochemical characteristics of RBD on its binding promiscuity. Our experiments from different binding assays, including human protein microarrays, indicated that both SARS-CoV-2 Wuhan and Omicron S proteins display binding promiscuity to unrelated proteins. However, the exact panels of off-targeted proteins preferentially recognized by both strains did not completely overlap. When analyzed in isolated form, the recombinant RBD from Wuhan strain manifested higher promiscuity as compared to RBD-δ and RBD-o. The changes in the pattern of promiscuity of the RBD can be well explained by physicochemical modification in its amino acid sequence. Thus, experiments with antibodies have demonstrated that structural dynamics of the antigen-binding site is one of the main factors determining the reactivity to unrelated antigens i.e. antibody polyreactivity ^57–62^. The structural dynamics of the RBD deduced by the signature of the activation thermodynamics can explain the tendency of the RBD-Wuhan for promiscuous interactions with unrelated to ACE2 proteins. A flexible binding site can accommodate through induced-fit to distinct molecular motifs. On the other hand, the display of extensive patches with positive charge on the molecular surface could explain the promiscuity of RBD-o, irrespectively of reduced structural dynamics. Indeed, studies with antibodies also provided evidence that the presence of positive patches in the antigen binding site is a strong driver of antigen-binding promiscuity ^63–66^. These facts could explain the observed protein binding promiscuity at the two ends of evolutionary trajectory of SARS-CoV-2. Importantly despite S protein retains its promiscuity, the different molecular mechanism of the off-target recognition conformational dynamics versus accumulation of positive charges provided explanation of the different set of recognized targets. These data might explain differences in cell tropism and infectivity of the different variants of SARS-CoV-2.

In conclusion of this study, we characterized the evolution of physicochemical mechanism of interaction of S protein of SARS-CoV-2 with its ligand. The obtained data revealed dramatic reconfiguration of binding mechanism in S protein from Omicron variant. These data could explain important functional traits of the virus such as cellular tropism, infectivity as well as immune evasion.

## Material and methods

### Proteins

Codon-optimized nucleotide fragments encoding stabilized versions of Wuhan and Omicron (BA.1) SARS-CoV-2 trimeric spike (HexaPro; S_6P) (S) ectodomains and receptor binding domains (RBD), followed by C-terminal tags (Hisx8-tag, Strep-tag, and AviTag) were synthesized and cloned into pcDNA3.1/Zeo(+) expression vector (Thermo Fisher Scientific). For SARS-CoV-2 RBD VOCs proteins, mutations (N501Y for the α variant; K417N, E484K and N501Y for the β variant; K471T, E484K and N501Y for the γ variant; L452R and T478K for the δ variant) were introduced using the QuickChange Site-Directed Mutagenesis kit (Agilent Technologies) following the manufacturer’s instructions. Glycoproteins were produced by transient transfection of exponentially growing Freestyle 293-F suspension cells (Thermo Fisher Scientific, Waltham, MA) using polyethylenimine (PEI) precipitation method as previously described ^67^. Proteins were purified from culture supernatants by high-performance chromatography using the Ni Sepharose® Excel Resin according to manufacturer’s instructions (GE Healthcare), dialyzed against PBS using Slide-A-Lyzer® dialysis cassettes (Thermo Fisher Scientific), quantified using NanoDrop 2000 instrument (Thermo Fisher Scientific), and controlled for purity by SDS-PAGE using NuPAGE 3-8% Tris-acetate gels (Sigma-Aldrich). AviTagged trimeric S and RBD proteins were biotinylated using the Enzymatic Protein Biotinylation Kit (Sigma-Aldrich).

### Evaluation of binding kinetics

Kinetics of interaction of SARS-CoV-2 S (Wuhan and Omicron variants) and of RBD (Wuhan, Alpha, Beta, Gamma, Delta and Omicron variants) proteins was evaluated by surface plasmon resonance-based biosensor system – Biacore 2000 (Cytiva Life Sciences, Biacore, Uppsala, Sweden). Recombinant ACE2 was covalently immobilized on CM5 sensor chips (Cytiva, Biacore) using amine-coupling kit (Cytiva, Biacore) according to the protocol recommended by the manufacturer. Briefly, ACE2 was diluted in 5 mM maleic acid (pH 4) to final concentration of 10 μg/ml and injected over sensor surfaces that were pre-activated by a mixture of 1-Ethyl-3-(3-dimethylaminopropyl)-carbodiimide and N-hydroxysuccinimide. For blocking the activated carboxyl groups thar were not engaged in interactions with the protein, the sensor surfaces were exposed to 1M solution of enthanolamine.HCl for 4 min. The solution of ACE2 was injected for contact times to achieve relatively low final coating densities of ∼150 and ∼450 RU. The coating of ACE2 and all kinetics measurements were performed by using as running buffer HBS-EP (10 mM HEPES pH 7.2; 150 mM NaCl; 3 mM EDTA, and 0.005 % Tween 20). The buffer was filtered through 0.22 µm filter and degassed under vacuum, before kinetics measurements.

To evaluate the binding kinetics of the interactions of S or RBD from different VOCs with ACE2, the viral proteins were serially diluted (two-fold each step) in HBS-EP. S-Wuhan and S-Omicron concentrations ranged from 10 to 0.156 nM and 5 to 0.0097 nM, respectively. The interaction RBD VOCs with immobilized ACE2 were measured after serial dilutions of 500–0.488 nM (WT, Alpha), 400, 200–0.39 nM (Beta, Gamma), 250–0.24 nM (Delta) and 31.25–0.12 nM (Omicron). The flow rate of samples and the running buffer during all interaction analyses was 30 µl/min. The association and dissociation phases of the binding of S proteins and different variants of RBD were monitored for 4 and 5 min, respectively. The sensor chip surfaces were regenerated by injection of solution of 4M guanidine hydrochloride for 30 sec. All kinetic measurements were performed at temperatures of 6, 12, 18, 24, 30 and 36 °C. The evaluation of the kinetic data was performed by BIAevaluation version 4.1.1 Software (Cytiva, Biacore) applying global and separate analyses models. For kinetic assessment of trimer S protein, a coupling density of ACE2 of ∼150 RU was used. In case of RBD variants the kinetics evaluation was performed using interactions obtained at binding density of the ligand of ∼450 RU.

### Evaluation of activation and equilibrium thermodynamics

Assessment of binding kinetics and changes in activation and equilibrium thermodynamic parameters of the interaction of RBD VOCs with ACE2 was performed according to a protocol described before ^57, 68, 69^. We applied Eyring’s analyses for evaluation of the activation thermodynamics of the interactions of RBD variants with ACE2. To this end the kinetic rate constants of association (k_a_) and dissociation (k_d_) obtained at distinct temperatures (6, 12, 18, 24, 30 and 36 °C) were used to construct Arrhenius plots. Slopes of the Arrhenius plots were calculated by applying linear regression analysis by using GraphPad Prism v.9 (GraphPad Prism Inc. USA) and were substituted in the equation -

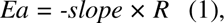

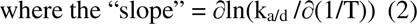

where *Ea* is the activation energy. The changes in activation enthalpy (ΔH^‡^), entropy (TΔS^‡^) and Gibbs free (ΔG^‡^), for association and dissociation phases were calculated by using the equations:

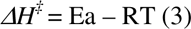

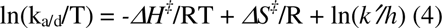

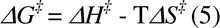

where T is the temperature in Kelvin degrees, *k′* is the Boltzman constant and *h* is the Planck’s constant.

The changes in thermodynamic parameters at equilibrim were calculated using the equations:

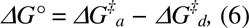

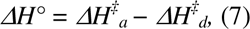

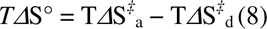

Activation and equilibrium thermodynamic parameters were determined at reference temperature of 24 °C (297.15 K).

### pH dependence of binding of RBD to ACE2

Ninety-six well polystyrene plates NUNC MaxiSorp™ (Thermo Fisher Scientific, Waltham, MA) were coated for 2 hours at room temperature with 10 μg/ml of ectodomain of human recombinant ACE2 in PBS. The plates were blocked for 1 hour by incubation at room temperature with solution of 1% of BSA in PBS. Next, site-specifically biotinylated RBD from SARS-CoV-2 Wuhan and VOCs (α; β; γ; δ, and ο) were diluted in buffers with increasing pH values in the range of 3-11 (0.5 pH unit increment) to final concentrations 0.625 μg/ml and 1 μg/ml, respectively. The buffer solutions maintaining pH range 3-7.5 were obtained by mixtures of different proportions of 0.1M citric acid and 0.2 M Na_2_HPO_4_. The buffers maintaining pH range 7.5-11 were based on mixtures of 0.1M Na_2_CO_3_ and 0.1M NaHCO_3_. All pH buffers contained 0.05 % Tween 20. The viral proteins diluted in buffers with different pH were incubated with immobilized ACE2 for 2 hours at room temperature. After the incubation and washing 5 × with PBS containing 0.05 % Tween 20 (T-PBS), the plates were incubated for 1 hour at room temperature with horseradish peroxidase-conjugated to streptavidin (R&D Systems), diluted 40× in T-PBS. Following thorough washing with T-PBS, the reactivity of RBD and S proteins was revealed by addition of peroxidase substrate, *o*-phenylenediamine dihydrochloride (Sigma-Adrich, St. Louis, MI). After stopping enzyme reaction by addition of 2M HCl, the absorbance was measured at 492 nm by using a microplate reader (Infinite 200 Pro, Tecan, Männedorf, Switzerland).

### Computational analyses of protein electrostatics

Coloring of the electrostatic charges were generated with the PyMOL built in APBS electrostatics plugin and visualized in PyMOL Molecular Graphics System (https://pymol.org/2/). The computational electrostatic calculations were performed with algorithms that were previously described in ^70, 71^. This software utilizes the mean force approach, written in Perl and C/C++ with Haskell extensions. It allows evaluation of the pH dependence of: (i) global electrostatic characteristics of proteins such as electrostatic free energy ΔEel (pH), protein net charge DZ(pH), electrostatic potential Φel, and value and direction of the dipole/electric moments vectors μd/μel(pH)) and local electrostatic potential [i(pH). All electrostatics calculations were done at a reference ionic strength of 0.1 M.e^2^.

### ELISA – Binding promiscuity of S protein

Ninety-six well polystyrene plates NUNC MaxiSorp™ (Thermo Fisher Scientific) were coated for 2 hours at room temperature with: human Factor IX; human serum albumin; human von Willebrand factor (all from LFB, Les Ulis, France); recombinant human Factor VIII (Helixate, Nextgen, Bayer, Leverkusen, Germany); human Factor B; human C3; human Factor H; human C1; human Factor I (all from Complement Technology, Tyler, TX); equine cytochrome C; bovine thyroglobulin; bovine histone III; human hemoglobin; chondroitin sulfate; keyhole limpet hemocyanin (all from Sigma-Aldrich); recombinant human nerve growth factor-β (R&D Systems, Minneapolis, MIN); LysM (from Enterococcus faecalis, kindly provided by Dr. Stephane Mesnage, University of Sheffield, UK), and recombinant human ACE2 (see above). All proteins were coated after dilution to 10 μg/ml in PBS. The chondroitin sulfate was coated at concentration of 500 μg/ml in PBS. The plates were blocked for 1 hour by incubation at 37 °C with solution of 0.25% in PBS. Next, site-specifically biotinylated S proteins from Wuhan and Omicron were diluted in T-PBS to 5 μg/ml and incubated for 90 minutes at room temperature. Next steps of ELISA assay were identical with the ones described above.

### ELISA – Inhibition of reactivity of S proteins by fetal calf serum

Ninety-six well polystyrene plates NUNC MaxiSorp™ (Thermo Fisher Scientific) were coated for 1 hour at 37 °C with recombinant human ACE2, with human recombinant nerve growth factor β (R&D Systems) and with bovine histone III (Sigma-Aldrich). All proteins were diluted to 10 μg/ml in PBS. After, the plates were blocked with 0.25 % solution of Tween 20 in PBS. S proteins from Wuhan and Omicron variants were diluted to final concentration of 1 μg/ml in T-PBS or in T-PBS containing fetal calf serum at two-fold dilutions in the range from 1/2 to 1/1052-fold diluted (50 to 0.097 %). The S proteins were pre-incubated for 10 min at room temperature and then transferred to plates with immobilized ACE2 and incubated for 1 hour at 37 °C. Next steps of ELISA assay were identical to the ones described with sole exception that instead of *o*-phenylenediamine dihydrochloride as a peroxidase substrate in these experiments was used TMB high sensitivity substrate solution (BioLegend, San Diego, CA) and absorbance was recorded at 450 nm.

### Conjugation of S proteins and RBD with fluorochrome for protein microarray and confocal microscopy analyses

RBDs (Wuhan, Delta and Omicron) and S proteins (Wuhan and Omicron) were covalently labeled with Alexa Fluor^TM^ 555. To this end, the proteins were first diluted to final concentration of 2 μM in buffer containing 2/3 PBS and 1/3 0.1M carbonate buffer with resulting pH of 8.3. Next, to the protein solutions was added Alexa Fluor^TM^ 555 NHS ester (Thermo-Fischer) with final concentration of 40 μM. Alexa Fluor^TM^ 555 NHS stock solution was prepared immediately before use by solubilizing the dye in DMSO (Sigma-Aldrich, >99.9 %) to stock concentration of 1.6 mM. The mixtures of the amine-reactive Alexa Fluor^TM^ 555 NHS ester with the proteins were incubated in dark with continuous shaking for 1 hour at room temperature. After, protein conjugates were separated from unconjugated fluorochrome by using PD-10 columns (Cytiva) pre-equilibrated with PBS. For estimation of efficacy of coupling of fluorochrome to RBDs and S proteins, the UV-visual absorbance spectra of conjugated proteins were recorded in the range 200–700 nm using Agilent Cary 100 absorbance spectrophotometer (Agilent Technologies, Santa Clara, CA).

### Protein microarrays

Reactivity of recombinant S proteins from Wuhan and VOC Omicron as well as of recombinant RBD (Wuhan, Delta and Omicron) to large set (>9000) of human proteins was assessed by using protein microarrays (ProtoArray^®^, Invitrogen, Thermo Fischer). First microarray chips were blocked by incubation for 1 hour at 4 °C with buffer containing: 50 mM HEPES, pH 7.5; 200 mM NaCl; 0.08 % Triton-X-100; 20 mM reduced glutathione; 1 mM dithiothreitol; 25 % glycerol, and 1 × synthetic block (Thermo Fischer). After washing 2 × 5 min with PBS containing 0.1 % Tween 20, the chips were incubated for 90 min at 4 °C with S proteins and RBDs labeled with Alexa Fluor^TM^ 555, diluted to 4.5 and 1 μg/ml, respectively in PBS containing 0.1 % Tween 20. After washing 2 × 5 min with PBS containing 0.1 % Tween 20, chips were soaked in deionized water, dried by centrifugation (200 × g) for 1 min. The fluorescence intensity was measured by using microarray scanner GenePix 4000B (Molecular Devices, San Jose, CA). The microarray chips were analyzed by using Spotxel software v. 1.7.7 (Sicasys, Heidelberg, Germany). The Z-scores for reactivity of SARS CoV-2 proteins to each human protein was determined by using ProtoArray Prospector v 5.2 software (Invitrogen, Thermo Fisher).

### Confocal microscopy

Human placental BeWo cells (2x10^5^ cells per well) were seeded on 24-well plates with a poly-L-lysine pre-coated coverslip in F-12 medium supplemented with 10% FCS. After 48 hours, cells were rinsed with PBS + CaCl_2_ + MgCl_2_ and incubated with 100 nM S proteins (Wuhan and Omicron) labeled with Alexa Fluor^TM^ 555 in presence or absence of 1% FCS at 4 °C during 30 min and after kept in a humidified incubator in 5% CO_2_ at 37 °C for 30 min. Cells were washed four times with ice-cold PBS + CaCl_2_ + MgCl_2_, fixed with 4% PFA for 20 min, and then permeabilized with 0.2% Triton-X-100 during 10 min. The cells were blocked for 35 min by incubation with solution of 1% of BSA in PBS and stained with 2 μg/mL Rabbit Anti-EEA1 antibody - Early Endosome Marker (ab2900, Abcam, Cambridge, UK) for 2 hours at room temperature, followed by 2 μg/mL Alexa Fluor^TM^ 488 goat anti-rabbit IgG (A11034, Invitrogen, Thermo Fisher) for 2 hours at room temperature. Nuclei were counterstained in blue with Hoechst 33342 dye (Invitrogen, Thermo Fisher). Cells were then washed, and coverslips were mounted in slides using fluoromount G (Invitrogen, Thermo Fisher). Images were acquired with a Zeiss LSM 710 inverted confocal microscope (Carl Zeiss AG, Germany). The quantification of the positive area (reflecting the spike interaction with the cells), normalized by the number of nuclei was measured using Halo Software, (Indica Lab, Albuquerque, NM). The positive area/cell for S proteins was normalized and the FCS-induced decrease was compared in all images of two independent experiments, for this, the average area/cell of S proteins was considered as 100 % and the values of each image in absence or presence of FCS of the two respective conditions (S-Wuhan or S-o) were divided by this denominator.

## Acknowledgments

This work was supported by Institut National de la Santé et de la Recherche Médicale (INSERM, France). J.D.D. is recipient of a grant from the European Research Council (Project CoBABATI ERC-StG-678905). A.R.R. is recipient of fellowship from an Innovative Training Network (ITN) funded by the European Union’s Horizon 2020 Programme under grant agreement No 859974, project EDUC8. A.Z. is supported by the Danish Council for Independent Research (DFF-1025-00015B).

